# Lower-limb Nonparametric Functional Muscle Network: Test-retest Reliability Analysis

**DOI:** 10.1101/2023.02.08.527765

**Authors:** Rory O’Keeffe, Jinghui Yang, Sarmad Mehrdad, Smita Rao, S. Farokh Atashzar

## Abstract

**Objective:** Functional muscle network analysis has attracted a great deal of interest in recent years, promising high sensitivity to changes of intermuscular synchronicity, studied mostly for healthy subjects and recently for patients living with neurological conditions (e.g., those caused by stroke). Despite the promising results, the between- and within-session reliability of the functional muscle network measures are yet to be established. Here, for the first time, we question and evaluate the test-retest reliability of non-parametric lower-limb functional muscle networks for controlled and lightly-controlled tasks, i.e., sit-to-stand, and over-the-ground walking, respectively, in healthy subjects.

**Method:** Fifteen subjects (eight females) were included over two sessions on two different days. The muscle activity was recorded using 14 surface electromyography (sEMG) sensors. The intraclass correlation coefficient (ICC) of the within-session and between-session trials was quantified for the various network metrics, including degree and weighted clustering coefficient. In order to compare with common classical sEMG measures, the reliabilities of the root mean square (RMS) of sEMG and the median frequency (MDF) of sEMG were also calculated.

**Results:** The ICC analysis revealed superior between-session reliability for muscle networks, with statistically significant differences when compared to classic measures.

**Conclusion and Significance:** This paper proposed that the topographical metrics generated from functional muscle network can be reliably used for multi-session observations securing high reliability for quantifying the distribution of synergistic intermuscular synchronicities of both controlled and lightly controlled lower limb tasks. In addition, the low number of sessions required by the topographical network metrics to reach reliable measurements indicates the potential as biomarkers during rehabilitation.

## I. Introduction

surface electromyography (sEMG) is a critical modality to measure muscular activities noninvasively on the skin.Technologies based on sEMG have been used for clinical decision-making and interfacing human neurophysiology with machine intelligence, for example, in powered prosthetic systems. The classic features of sEMG, such as root mean square (RMS), have demonstrated acceptable within-session test-retest reliability with an intraclass correlation coefficient (ICC) > 0.75 during some everyday dynamic tasks, such as walking or cycling [1], [2], [3], [4]. On the other hand, between-session reliability of the classic features has shown weaker results. Acceptable between-session reliability (ICC > 0.75) has been demonstrated by the RMS and mean frequency for isometric tasks by some studies [5], [6] but not by others [7]. For more complex movements, the sEMG may vary between sessions [8], [9], and thus between-session reliability for everyday dynamic tasks is not supported in the literature [9]. This lack of reliability can significantly challenge the use of this modality for tracking changes in the neurophysiological functionalities that may be induced by rehabilitation, exercise, degenerative conditions, and aging. High between-session reliability would be needed for generating any clinical biomarker [10].

This variability hinders efforts to use sEMG metrics as effective biomarkers to track muscular activity across impairments and rehabilitation. In this paper, we hypothesize that by going beyond individual muscle activation and by focusing on the underlying distribution of the neural drive, and decoding patterns of information sharing between muscles, the needed ICC would be realized for both between-session and within-session reliability analyses. To do this, we will conduct the ICC analysis in this paper to assess reliability between and within sessions when compared with classic spectrotemporal measures. In addition, we will utilize the Spearman-Brown prophecy formula to understand how many measurements would be needed to reach a consistent result [11], [12].

Recently, the concept of functional muscle network has been proposed to model systematic activation synchronization between skeletal muscles. Muscle network has provided promising results in discriminating subtle motor differences [13], [14]. Our recent studies have shed light on the clinical benefits of various formulations of muscle networks. In this regard (a) we have recently shown that nonlinear informationtheory-based temporal-domain functional muscle networks of skeletal muscles provide excellent conformity for the active muscles across stroke subjects and show high sensitivity to sensorimotor integration [15]; (b) we have shown that linear frequency-domain functional muscle network of perilaryngeal muscles can discriminate subtle changes in vocal tasks and be a potential biomarker of vocal hyperfunction [16], [17]; (c) we have also shown that non-parametric muscle network can be a sensitive biomarker of lower-limb fatigue with superior sensitivity when compared with conventional spectrotemporal biomarkers of fatigue [18]. As the next significant step, in this paper and for the first time, we investigate whether non-parametric functional muscle networks can secure test-retest reliability for within- and between-session settings during lightly-controlled lower limb tasks, which would be needed for any biomarker tracking the performance.

It should be noted that the muscle network is composed of a set of nodes (muscles) and edges (the connectivity between the two nodes) [19], [20]. The edges can be calculated using linear or nonlinear models and can be processed in the frequency or time domain [21], [22], [23]. Forming the muscle network facilitates the employment of network theory and extraction of topographical graph features that explain the network of muscles rather than individual muscles. It explains the distribution of synergistic actions and patterns of information sharing between muscles. Examples of such features are the network Degree and weighted clustering coefficient (WCC), which respectively measure functional integration and segregation, respectively [24], [25]. The implementation of the network-theoretic topographical measures may increase reliability as these measures focus on the network structure (less sensitive to individual channel noises) rather than the specific pairwise connectivities.

In this paper, we focus on the sit-to-stand and walking tasks as they are of great importance in everyday life and in clinical and rehabilitation contexts. Previous studies have demonstrated that sit-to-stand is a useful test of composite lower extremity muscle strength [26], [27]. Due to the essential nature of the sit-to-stand task in everyday life, metrics relating to the task are analyzed in clinical settings [28], [29], [30]. During a walking task, the lower limb muscles activate to a sub-maximal level in a coordinated manner [31], [32], [33]. Walking tasks such as timed up and go and other standardized tests are used to track progress for patients during rehabilitation from a variety of conditions such as skeletal disorders, heart disease, stroke, cognitive impairments, and others [34], [35], [36], [37]. Due to the high relevance of both tasks in a clinical context, results from this work could shed light on the most reliable electromyographic biomarkers for tracking progress during rehabilitation.

Contributions of this paper are explained as follows. In this study, we investigate the within- and between-session reliability of muscle network metrics during non-demanding and loosely-controlled tasks, such as sit-to-stand and walking. Our primary hypothesis is that the functional muscle network, which takes into account the coordination and interaction of multiple muscle groups, would exhibit higher between-session reliability than traditional metrics such as RMS and median frequency (MDF) derived from sEMG measurements. To test this hypothesis, we calculate the intraclass correlation coefficient and conduct the Spearman-Brown prophecy analysis. Our findings suggest that the functional muscle network metrics show stronger between-session reliability for both walking and sit-to-stand tasks compared to traditional metrics. These results have important implications for the use of muscle network metrics in tracking progress during rehabilitation. This paper suggests that muscle networks provide a more comprehensive and reliable representation of muscle function compared to traditional metrics and secure test-retest reliability performance for the lower limb.

## II. Methods

Fifteen asymptomatic subjects (eight females) with an age of 25.9 ± 3.6 (mean ± standard deviation (SD)) years and a BMI of 22.8 ± 3.1 (mean ± SD) *kg/m*^2^ participated in the study. The study was approved by the Institutional Review Board of New York University, and subjects provided their written consent prior to the start of the experiment. Subjects did not report any lower-limb injury, pain, or impairment at the time of the experiment.

### A. Experiment Procedure

Subjects performed 30 seconds of sit-to-stand and 30 seconds of walking in two sessions that were apart at least one week from each other (Fig. 1a). For the sit-to-stand task, subjects were instructed to repeatedly sit and stand up from a fixed chair with arms folded across their chest at a comfortable pace. The number of repetitions completed by each subject in each session is summarized in Table I. Similarly, subjects walked over the ground in a straight line at their comfortable pace for 30 seconds. We did not control the pace or provide any instructions as to how to turn when subjects reached the end of the path. The number of strides during walking in each session is summarized in Table II.

**TABLE I.** Sit-to-stand Repetitions for session 1 and 2

| Subject | Session 1 | Session 2 |
| --- | --- | --- |
| 1 | 21 | 22 |
| 2 | 20 | 21 |
| 3 | 12 | 11 |
| 4 | 12 | 10 |
| 5 | 25 | 29 |
| 6 | 7 | 7 |
| 7 | 10 | 10 |
| 8 | 10 | 8 |
| 9 | 8 | 8 |
| 10 | 11 | 10 |
| 11 | 12 | 15 |
| 12 | 8 | 8 |
| 13 | 9 | 10 |
| 14 | 10 | 9 |
| 15 | 8 | 8 |
| Mean | 12.20 | 12.40 |
| Std | 5.40 | 6.50 |

**TABLE II.** Walking stride counts for sessions 1 and 2

| Subject | Session 1 | Session 2 |
| --- | --- | --- |
| 1 | 11 | 11 |
| 2 | 14 | 14 |
| 3 | 16 | 16 |
| 4 | 12 | 13 |
| 5 | 12 | 11 |
| 6 | 11 | 11 |
| 7 | 10 | 10 |
| 8 | 11 | 12 |
| 9 | 13 | 12 |
| 10 | 12 | 10 |
| 11 | 11 | 11 |
| 12 | 12 | 13 |
| 13 | 13 | 14 |
| 14 | 13 | 12 |
| Mean | 12.21 | 12.14 |
| Std | 1.53 | 1.70 |

**Fig. 1.**
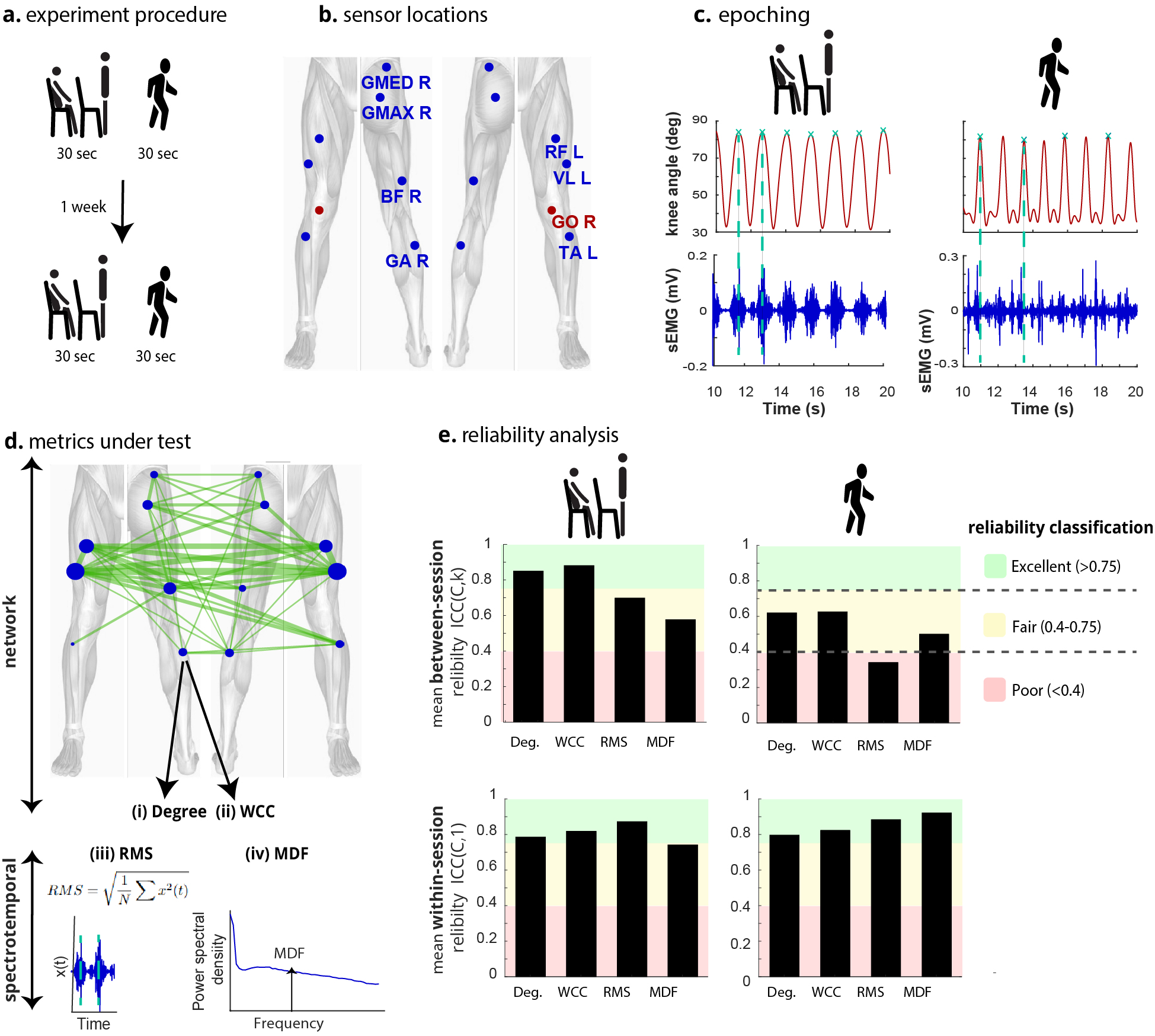
Test-retest reliability of the network and spectrotemporal metrics for everyday tasks. **a**. Subjects completed two sessions of the 30s of sit-to-stand and walking at a natural pace. **b**. Seven bipolar sEMG sensors were placed bilaterally. GMED = Gluteus Medius, GMAX = Gluteus Maximus, BF = Biceps Femoris, GA = Gastrocnemius, RF = Rectus Femoris, VL = Vastus Lateralis, TA = Tibialis Anterior. Goniometers were placed bilaterally to measure the knee angle. **c**. Epoch limits were defined using the goniometer, and all metrics were computed epoch-wise **d**. The functional muscle network was constructed by quantifying the absolute Spearman power correlation (|*ρ*_*ij*_|) between muscle pairs. (i) Degree and (ii) weighted clustering coefficient (WCC) were used to quantify the network connectivity at each muscle (node). (iii) Root mean square (RMS) and (iv) median frequency (MDF) were quantified as spectrotemporal measures. (iv) The MDF divides the area under the power spectrum in two. **e**. The main difference between network and spectrotemporal metrics was in between-session ICC. For sit-to-stand, the mean between-session ICC was excellent for the network metrics but was fair for RMS and MDF. For walking, between-session ICC was fair for the network metrics, poor for RMS, and fair for MDF. The within-session ICC was excellent for all metrics.

The sEMG signals were recorded from seven lower-limb muscles on each side: 1) Rectus Femoris (RF), 2) Vastus Lateralis (VL), 3) Biceps Femoris (BF), 4) Gluteus Medius (GMED), 5) Gluteus Maximus (GMAX), 6) Gastrocnemius (GO) and 7) Tibialis Anterior (TA), using wireless Trigno system (Delsys Inc, Natick, MA) at 1259 Hz. We used two goniometers (Biometrics Ltd, UK) to record knee angles at 296 Hz throughout the tests. The placement was made according to SENIAM guidelines, and the skin area was properly wiped using alcohol pads. The data was exported to the MATLAB environment (R2020b, MathWorks Inc., Natick, MA) for processing. The first and last one second of each trial were removed. The tasks were divided into epochs for calculating the network metrics and spectrotemporal measurements. The peaks of the right knee angle delineated the epochs for sit-to-stand and walking (Fig. 1c). The start and end time of the *i*^*th*^ epoch were the respective times at which the *i*^*th*^ and (*i* + 1)^*th*^ peaks occurred. For walking, every other peak of the right knee angle was used to delineate the gait cycle epochs.

### B. Muscle Network

In this paper, the non-parametric Spearman power correlation network was constructed for each epoch of the sEMG signals that were bandpass-filtered between 20 and 50 Hz [18]. In short, the Spearman power correlation quantifies the monotonic trends between the power of two signals, even if they are not linearly related. The Spearman correlation coefficient ranges from -1 to 1, which indicates ideal monotonic reverse relation to ideal monotonic relation, respectively. The absolute Spearman power correlation (|*ρ*_*ij*_|) was used to quantify the monotonic relationship and hence connectivity between muscle *I* and muscle *J*. The network was constructed using the pairwise absolute Spearman power correlation |*ρ*_*ij*_| where muscles act as nodes and the connectivity values act as edges between the nodes.

The degree of each node, *D*_*i*_, is then defined as the average of all edges connected to the node. Assuming the muscle network as an adjacency matrix *A, D*_*i*_ is defined as:

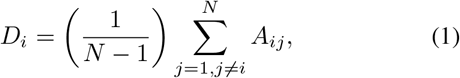

where *N* is the number of nodes. Mean network degree, is the mean of all nodes’ degrees:

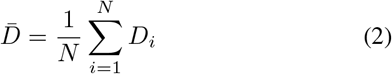

where *N* is the number of nodes (i.e. muscles).

The weighted clustering coefficient (*WCC*_*i*_) gives a relative measure of how well node *i* is connected to its neighbors (*A*_*ij*_, *A*_*ik*_) while also accounting for the neigbors’ interconnection (*A*_*jk*_). *WCC*_*i*_ can be defined as:

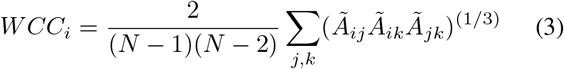

*WCC*_*i*_ = 0 represents an isolated node and *WCC*_*i*_ = 1 represents a fully connected node to its neighbors. Mean network WCC, is the mean of all nodes’ WCCs:

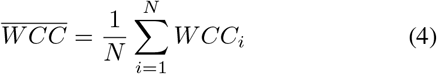

### C. Conventional Spectrotemporal Measurements

To compare the proposed network results to the existing attempted biomarker methods, RMS and MDF were quantified for each epoch with respect to particular muscles. The sEMG bandwidth of 20-200 Hz was considered for RMS and MDF. The signal was band-pass filtered between 20 and 200 Hz with a zero-phase 4th order Butterworth filter in addition to applying zero-phase 4th order Butterworth notch filters (half-width = 2.5 Hz) at multiples of 60 Hz. RMS and MDF were computed using the unrectified filtered signal. The RMS of a muscle signal *x*(*t*) is defined as:

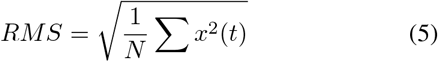

where *N* is the number of time points. The MDF is defined as that frequency that divides the area under the power spectrum density curve in two; hence it indicates the power spectrum shape, while the median PSD indicates the overall magnitude.

### D. Test-retest reliability

Intraclass correlations were evaluated for both between- and within-session test-retest reliability analysis [38], [39]. We used the SPSS software (version 28, IBM Corp, Armonk, NY) to compute the ICCs. Network and spectrotemporal metrics were quantified epoch-wise during each task.

To measure the between-session reliability of each metric, the intraclass correlation coefficient was calculated with the fixed column and random row effects model, average score (*ICC*(*C, k*)) [39], [40]. *ICC*(*C, k*) was quantified from a data matrix in the standard format presented by McGraw and colleagues with the subjects on the rows and the sessions on the columns [39], as illustrated in Table III. *ICC*(*C, k*) indicates the level of consistency between the two sessions, and can be interpreted as the reliability of the average measurement across sessions [38], [39]. Each element of the data matrix *X* represents the mean metric across epochs for the *i*^*th*^ subject in a given session.

**TABLE III.** Data matrix format for the calculation of between-session reliability with *ICC*(*C, k*).

| Subject | Session |  |
| --- | --- | --- |
|  | 1 | 2 |
| 1 | $X_{11}$ | $X_{12}$ |
| ... | ... | ... |
| i | $X_{i1}$ | $X_{i2}$ |
| 15 | $X_{151}$ | $X_{152}$ |

For the within-session analysis, the ICC was calculated with the fixed column and random row effects model, single score (*ICC*(*C*, 1)). *ICC*(*C*, 1) was quantified from a data matrix in the standard format with the subjects on the rows and the epochs on the columns as illustrated in Table IV. *ICC*(*C*, 1) indicates the level of consistency between the epochs and can be interpreted as the reliability of a single epoch measurement (4) [38], [39]. Each element of the data matrix *Y* represents a metric value for the *i*^*th*^ subject in the *j*^*th*^ epoch. The minimum number of epochs across all subjects was chosen as the number of observations (*m*) for each subject. The mean *ICC*(*C*, 1) value across both sessions was used to estimate the withinsession reliability.

**TABLE IV.** DATA MATRIX FORMAT FOR THE CALCULATION OF THE WITHIN-SESSION RELIABILITY WITH *ICC*(*C*, 1)

| Subject | Epoch |  |  |  |
| --- | --- | --- | --- | --- |
|  | 1 | ... | j | m |
| 1 | $Y_{11}$ | ... | $Y_{1j}$ | $Y_{1m}$ |
| ... | ... | ... | ... | ... |
| i | $Y_{i1}$ | ... | $Y_{ij}$ | $Y_{im}$ |
| 15 | $Y_{151}$ | ... | $Y_{15j}$ | $Y_{15m}$ |

ICC values calculated as negative were set to 0. The *ICC*(*C, k*) and *ICC*(*C*, 1) of a given metric were computed for each muscle, and the results are summarized in Fig. 1e where the mean ICCs of each metric across muscles are shown. In the literature, the reliability of sEMG metrics can be interpreted from the ICCs using Fleiss’ scale [41], [3] as follows: poor (*ICC* < 0.4), fair (0.4 < *ICC* < 0.75) or excellent (*ICC* > 0.75). This interpretation is illustrated in Fig. 1e and continued throughout the paper.

### E. Spearman-Brown Analysis

In the rest of this section, based on the observed behavior, we evaluate how many sessions are needed to statistically claim the excellent ICC (*ICC >* 0.75). It has been shown that increasing the number of measurements enhances the average-score reliability [38], [39]. Given the current reliability *r*, the factor *n* by which the number of measurements needs to increase so that the desired average-score reliability *r*^*∗*^ would be attained can be calculated using the Spearman-Brown prophecy formula [42], [43], [12].

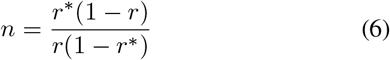

For excellent between-session reliability, we would require an *ICC*(*C, k*) ≥ 0.75. Hence:

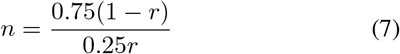

The hypothetical number of sessions that would be required to attain excellent reliability (*N*_*S*_) was calculated by multiplying *n* by the current number of sessions (2) and rounding up.

Hence:

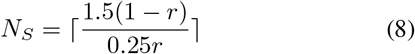

where *r* is the current between-session reliability. *N*_*S*_ is shown for sit-to-stand and walking in Table V and Table VI respectively with values provided for all metrics. Note that if *r* = 0, then *N*_*S*_ = ∞.

**TABLE V.** **S****it-to-stand:** Number of sessions needed to achieve excellent between-session reliability (*N*_*S*_)

| Muscle | Metric<br>Degree | WCC | RMS | MDF |
| --- | --- | --- | --- | --- |
| RF L | 1 | 1 | 2 | 2 |
| VL L | 1 | 1 | 1 | 1 |
| BF L | 1 | 1 | 58 | 3 |
| GMED L | 3 | 2 | 42 | 13 |
| GMAX L | 2 | 1 | 2 | ∞ |
| GA L | 2 | 2 | 1 | ∞ |
| TA L | 2 | 1 | 2 | 8 |
| RF R | 2 | 1 | 2 | 2 |
| VL R | 1 | 1 | 2 | 2 |
| BF R | 1 | 1 | 3 | 2 |
| GMED R | 2 | 1 | 2 | 2 |
| GMAX R | 1 | 1 | 2 | 7 |
| GA R | 3 | 3 | 5 | 5 |
| TA R | 2 | 1 | 3 | 5 |
| Mean L | 1.71 | 1.29 | 15.43 | N/A |
| Mean R | 1.71 | 1.29 | 2.71 | 3.57 |
| Mean | 1.71 | 1.29 | 9.07 | N/A |

**TABLE VI.** TEXTBFWalking: Number of sessions needed to achieve excellent between-session reliability (*N*_*S*_)

| Muscle | Metric<br>Degree | WCC | RMS | MDF |
| --- | --- | --- | --- | --- |
| RF L | 1 | 1 | 4 | 4 |
| VL L | 2 | 2 | 3 | 23 |
| BF L | 3 | 3 | 34 | 3 |
| GMED L | 62 | 24 | ∞ | 8 |
| GMAX L | 6 | 6 | 8 | 31 |
| GA L | 11 | 8 | ∞ | 5 |
| TA L | 4 | 5 | ∞ | 15 |
| RF R | 2 | 3 | 2 | 8 |
| VL R | 1 | 2 | 23 | 2 |
| BF R | 4 | 4 | 2 | 7 |
| GMED R | 4 | 3 | ∞ | 4 |
| GMAX R | 20 | 11 | 31 | 254 |
| GA R | 2 | 3 | 5 | 7 |
| TA R | 5 | 3 | 305 | 2 |
| Mean L | 12.71 | 7 | N/A | 12.71 |
| Mean R | 5.43 | 4.14 | N/A | 40.57 |
| Mean | 9.07 | 5.57 | N/A | 26.64 |

## III. Results

With respect to the sit-to-stand task, the network metrics (degree and WCC) demonstrated both excellent between- and within-session reliability. For the between-session analysis, the average reliability of both degree and WCC across left-sided, right-sided, and all muscles was excellent (mean *ICC*(*C, k*) > 0.75, Figs. 1e, 2a). The degree of 12/14 muscles and the WCC of 13/14 muscles displayed excellent between-session reliability (*ICC*(*C, k*) > 0.75, Fig. 2a). For the within-session analysis, the average reliability of each network metric across left-sided, right-sided, and all muscles was excellent (mean *ICC*(*C*, 1) > 0.75, Figs. 1e, 2b). The degree of 9/14 muscles and the WCC of 12/14 muscles displayed excellent within-session reliability (*ICC*(*C*, 1) > 0.75, Fig. 2b). Overall, the network degree and WCC for VL, BF, RF, and gluteus muscles (GMED and GMAX) showed excellent between- and within-session reliability for the sit-to-stand task. The ankle dorsi and plantar flexors (TA and GA) had the minimum ICC (both within and between sessions) among all muscles, yet their reliability was within the statistically fair range (degree of GA L: *ICC*(*C*, 1) ∼0.5, Fig. 2b).

**Fig. 2.**
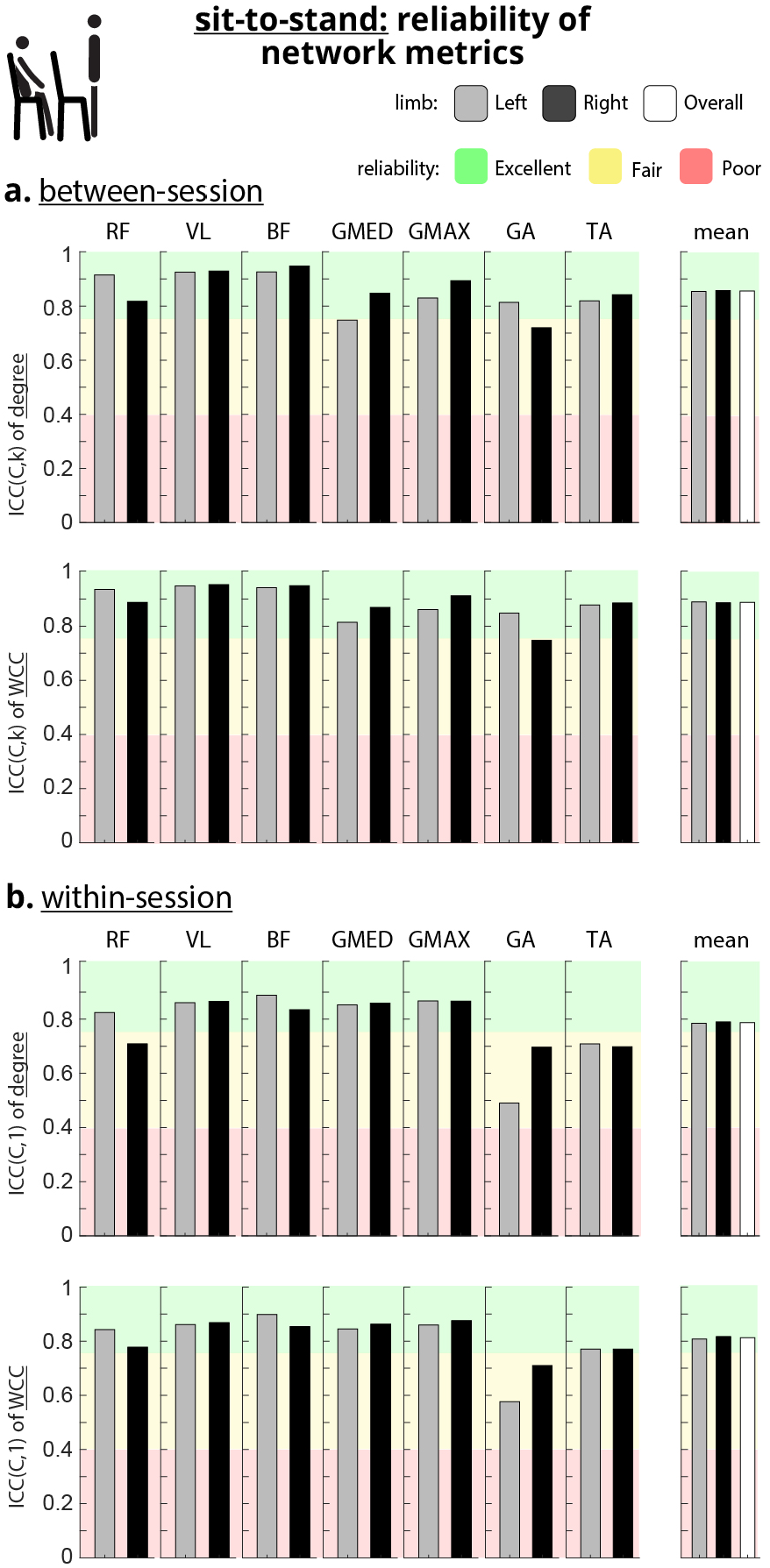
Reliability of network metrics for the sit-to-stand task. **a**. Betweensession (*ICC*(*C, k*)) reliability of the degree (top) and WCC of all fourteen muscles. The mean between-session reliability of each network metric across left-sided, right-sided, and all muscles was excellent. **b**. Within-session (*ICC*(*C*, 1)) reliability of the degree (top) and WCC of all fourteen muscles.

On the other hand, the spectrotemporal metrics (RMS and MDF) demonstrated only fair between-session reliability and mixed performance for the within-session reliability with respect to sit-to-stand. For the between-session analysis, the average reliability of both RMS and MDF across all muscles was fair (0.75 > mean *ICC*(*C, k*) > 0.4, Figs. 1e, 3a). The RMS of 8/14 muscles and the MDF of 6/14 muscles displayed excellent between-session reliability (*ICC*(*C, k*) > 0.75, Fig. 3a). However, the RMS of 4/14 and the MDF of 5/14 muscles showed fair between-session reliability (0.75 > *ICC*(*C, k*) > 0.4, Fig. 3a). The RMS of 2/14 muscles, including BF L and GMED L, and the MDF of 3/14 muscles, including GMED L, GMAX L, and GA L, showed poor between-session reliability (*ICC*(*C, k*) < 0.4, Fig. 3a). For the within-session analysis, the average reliability of RMS across left-sided, right-sided, and all muscles was excellent (mean *ICC*(*C*, 1) > 0.75, Figs. 1e, 3b), but the average reliability of MDF across all muscles was fair (0.75 > mean *ICC*(*C*, 1) > 0.4, Fig. 3b). The RMS of 11/14 and the MDF of 7/14 muscles displayed excellent within-session reliability (*ICC*(*C*, 1) > 0.75, Fig. 3b).

**Fig. 3.**
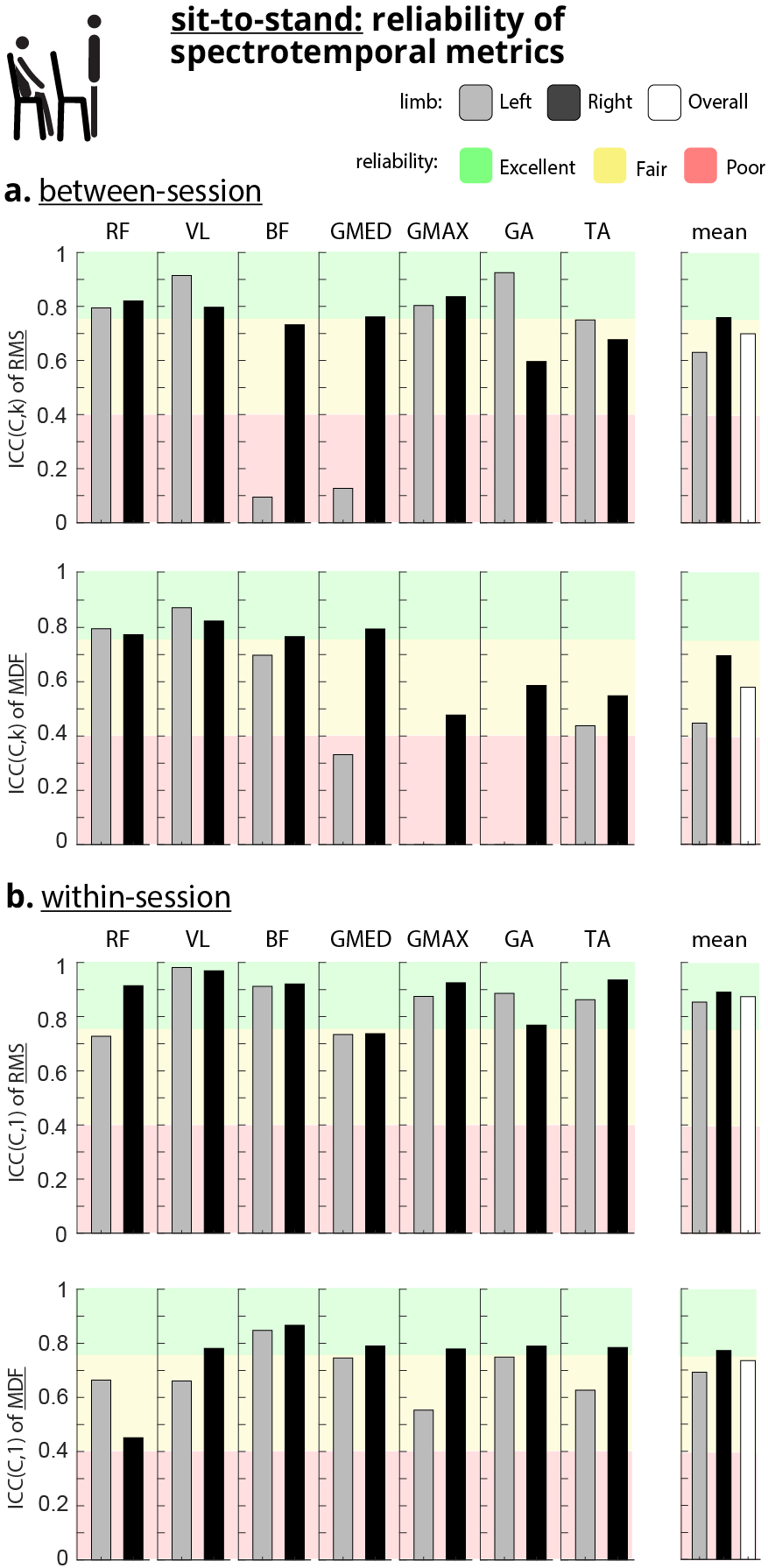
Reliability of spectrotemporal measurements for the sit-to-stand task. **a**. Between-session (*ICC*(*C, k*)) reliability of the RMS (top) and MDF of all fourteen muscles. The mean between-session reliability of each spectrotemporal measurement across all muscles was fair. **b**. Within-session (*ICC*(*C*, 1)) reliability of the RMS (top) and MDF of all fourteen muscles.

With respect to the walking task, both network metrics showed fair between- and excellent within-session reliability. For the between-session analysis, the average reliability of each network metric across left-sided, right-sided and all muscles was fair (mean *ICC*(*C, k*) > 0.4, Figs. 1e, 4a). The degree of 5/14 muscles and the WCC of 3/14 muscles displayed excellent between-session reliability (*ICC*(*C, k*) > 0.75, Fig. 4a). For the within-session analysis, the average reliability of each network metric across left-sided, right-sided and all muscles was excellent (mean *ICC*(*C, k*) > 0.75, Figs. 1e, 4b). The degree of 13/14 muscles and the WCC of 13/14 muscles displayed excellent within-session reliability (*ICC*(*C, k*) > 0.75, Fig. 4b).

**Fig. 4.**
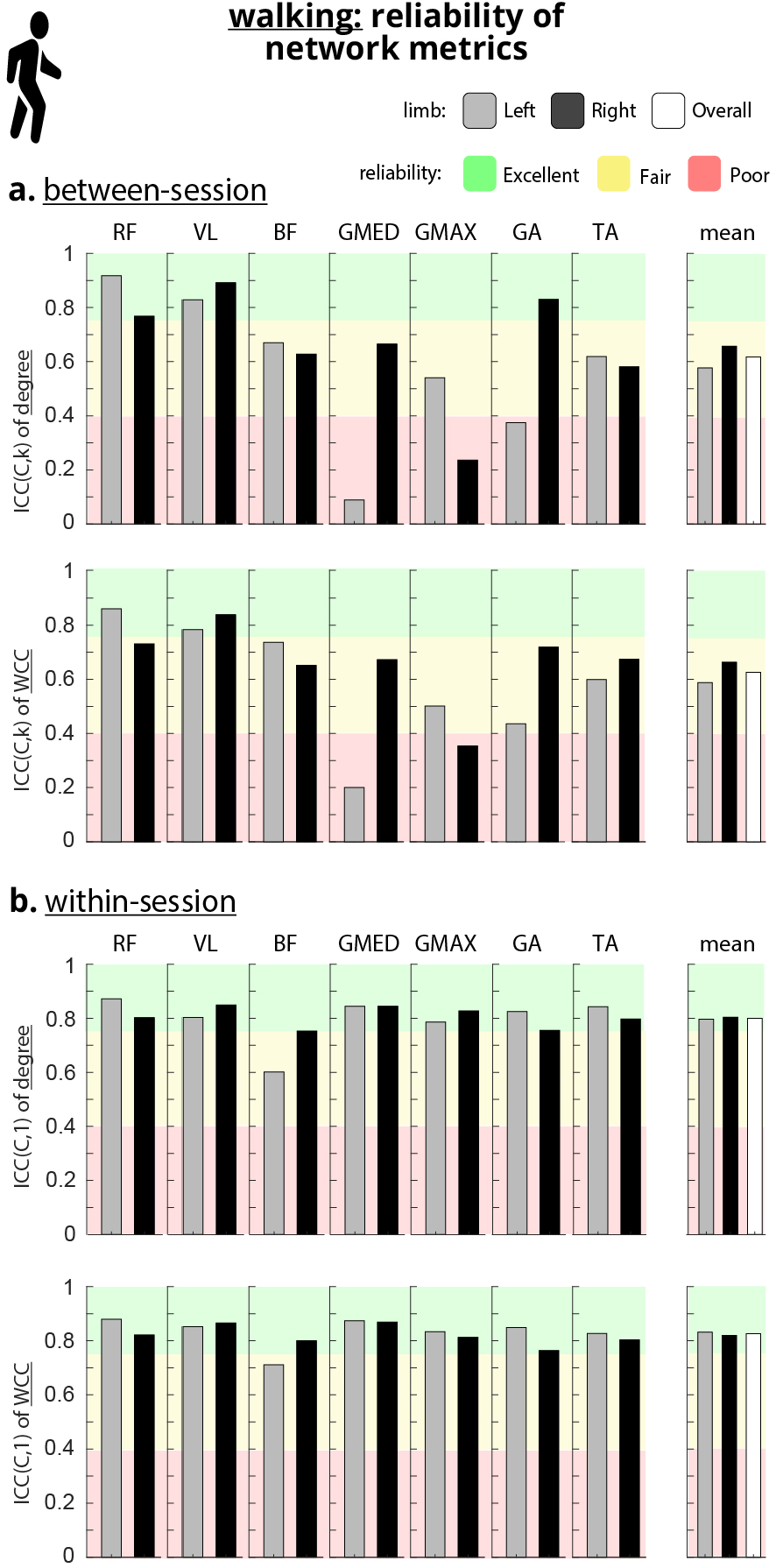
Reliability of network metrics for walking. **a**. Between-session (*ICC*(*C, k*)) reliability of the degree and WCC of all fourteen muscles. The degree of each quadricep muscle (RF and VL) had excellent between-session reliability. **b**. Within-session (*ICC*(*C*, 1)) reliability of the degree and WCC of all fourteen muscles. The mean within-session reliability of each network metric across left-sided, right-sided, and all muscles was excellent.

On the other hand, spectrotemporal metrics displayed poor to fair between-session reliability while also showing excellent within-session reliability with respect to walking. For the between-session analysis, the average reliability of RMS was poor (mean *ICC*(*C, k*) < 0.4, Figs. 1e, 5a) while the average reliability of MDF was fair (0.75 > mean *ICC*(*C, k*) > 0.4, Figs. 1e, 5a). Only 2/14 muscles displayed excellent between-session reliability for each of RMS and MDF (*ICC*(*C, k*) > 0.75, Fig. 5a). However, the average within-session reliability of each spectrotemporal metric across left-sided, right-sided, and all muscles was excellent (mean *ICC*(*C, k*) > 0.75, Fig. 1e, Fig. 5b). The RMS of 13/14 and the MDF of 14/14 muscles displayed excellent within-session reliability (*ICC*(*C*, 1) > 0.75, Fig. 5b).

**Fig. 5.**
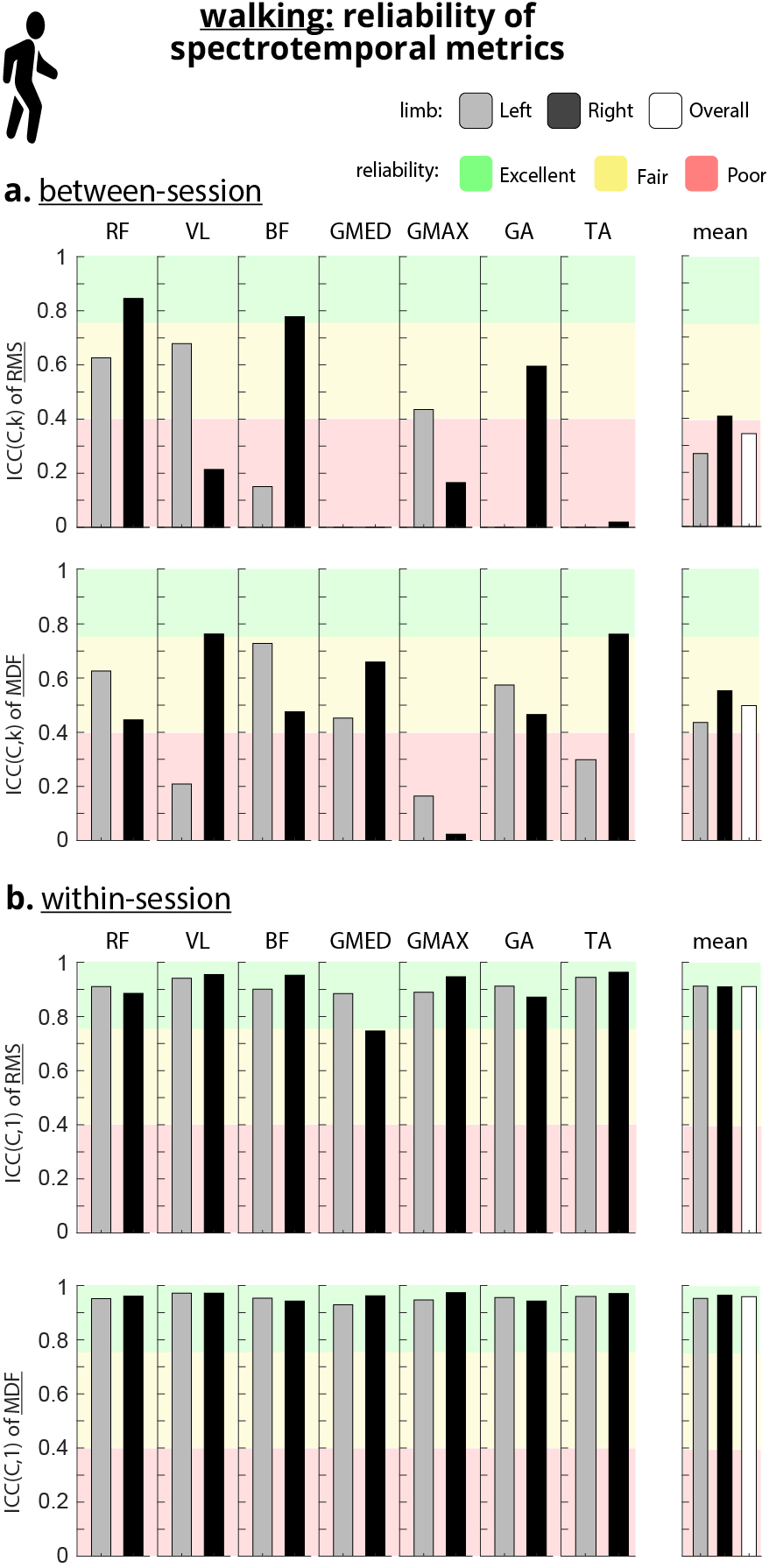
Reliability of spectrotemporal metrics for the walking task. **a**. Between-session (*ICC*(*C, k*)) reliability of the RMS and MDF of all fourteen muscles. **b**. Within-session (*ICC*(*C*, 1)) reliability of the RMS and MDF of all fourteen muscles. The mean within-session reliability of each spectrotemporal metric across left-sided, right-sided and all muscles was excellent.

Using the Spearman-Brown prophecy formula and the ICC values, the hypothetical number of sessions needed for excellent reliability (*N*_*S*_) was determined. For the sit-to-stand task, the network metrics required fewer sessions to attain excellent reliability than the spectrotemporal metrics (Table V). The mean *N*_*S*_ across all muscles was 1.71, 1.29, 9.07, and *N/A* for the degree, WCC, RMS, and MDF, respectively. Since MDF had *N*_*S*_ = ∞ for some muscles, the mean could not be calculated. The number of muscles that required only a single session for excellent reliability was 6/14, 11/14, 2/14, and 1/14 muscles for the degree, WCC, RMS, and MDF, respectively. The metric with the lowest *N*_*S*_ (colored green) was most frequently WCC which had the lowest *N*_*S*_ for 12/14 muscles and the lowest mean *N*_*S*_ across left, right, and all muscles. For the walking task, the network metrics required fewer sessions to attain excellent between-session reliability than the spectrotemporal metrics (Table VI). The mean *N*_*S*_ across all muscles was 9.07, 5.57, *N/A*, and 26.64 for the degree, WCC, RMS, and MDF, respectively. Since RMS had *N*_*S*_ = ∞ for some muscles, the mean could not be calculated. The degree of RF L and VL-R and the WCC of RF L would attain excellent reliability from a single session of walking, while the spectrotemporal metrics of all muscles need at least two sessions to attain such reliability. The metric with the lowest *N*_*S*_ (colored green) was most frequently degree which had the lowest *N*_*S*_ for 8/14 muscles, but WCC had the lowest mean *N*_*S*_ across left, right, and all muscles.

## IV. Discussion

This is the first study to evaluate the between-session reliability of the functional muscle network. We showed that the muscle network metrics have excellent within- and between-session reliability. The results demonstrated superior between-session reliability of the network compared to spectrotemporal metrics (Figs. 4a, 5a) and suggest that network metrics have more consistent behavior across sessions of a dynamic task, and hence have an increased likelihood to detect meaningful neurophysiological changes. In this regard, it should be highlighted that the between-session reliability of network metrics significantly outperforms the spectrotemporal counterparts for both sit-to-stand and walking (Fig. 1e), which supports our central hypothesis.

The within-session reliability of network metrics was on the same level as that of spectrotemporal counterparts for both sit-to-stand and walking (Fig. 1e). The significantly higher between-session reliability of the topographical features of the muscle network, besides excellent within-session reliability, supports the use of it as a biomarker when it detects and tracks changes in human neurophysiology over various days of assessments or interventions. In other words, if and when changes are observed, it can be reliably considered as the result of the understudied condition/procedure rather than natural random changes.

For example, the results of this study support the extrapolation of recent successes of muscle networks in detecting neurophysiological changes caused by the following conditions to go beyond observational assessment and approach development of novel biomarkers: (a) stroke and sensory stimulation (see our recent work [15]), (b) fatigue in healthy subjects (see our recent work [18]), or (c) underlying muscle hypertension and various functional tasks (see our recent work [17], [16]). At the node/muscle level, the excellent reliability for the network metrics of targeted muscles during the dynamic tasks (for example, VL bilaterally, Fig. 4a) shows that the muscles that are specifically the main contributors to the tasks would also indicate robust network measures both within and between sessions.

It should be highlighted that using a similar formulation of muscle network as the one used in this current paper, in our recent observational study, we investigated the behavior of network degree and weighted clustering coefficient when compared to classic spectrotemporal measures [18]. In [18] we showed that the network metrics can detect significant changes before to after fatigue protocol in a consistent manner, at the group, individual, and muscle levels while spectrotemporal metrics failed to detect consistently. The superior between-session reliability of the network over spectrotemporal metrics displayed here (Figs. 2a, 3a) supports the observations made for the use of muscle network to detect fatigue and would link the changes to the intervention (here is fatigue) rather than natural variations between two distinct measurements, further underlining the importance and significance of the made observations.

The within-session results affirm the excellent reliability of the network metrics during the dynamic tasks. Different from between-session reliability, in which the spectrotemporal measures fail to secure the needed robustness, in this paper, for the first time, we showed that both topographical measures from the network and the spectrotemporal measures secure high within-session reliability. The high within-session reliability would likely facilitate the detection of different modes of coordination within the same trial and would support the extrapolation of results from studies such as [13].

In contrast to network metrics, spectrotemporal metrics did not demonstrate robustness to day-to-day variabilities. In this regard, the corresponding average between-session reliability of spectrotemporal metrics was either fair or poor (Fig. 1e).

Indeed, very poor between-session reliability (*ICC*(*C, k*) < 0.2, Fig. 3a) was observed for spectrotemporal metrics for four left-sided muscles for the sit-to-stand task, and this could be due to higher variability on the subjects’ non-dominant side (14 out of 15 subjects were right-footed). The non-dominant side is known to have a weaker motor control [44], [45] and the sEMG characteristics of non-dominant lower-limb muscles can be distinguished from those of the dominant side [46], [47]. To answer this question, there would be a need for a follow-up study (out of the scope of this paper) to compare the effect of the dominant side on the reliability of spectrotemporal measures in comparison with that of network measures.

As established above, the presented results suggest that the spectrotemporal metrics are more susceptible to myographic variabilities. The higher variability and thus lower reliability of spectrotemporal metrics in both tasks (Figs. 2a, 3a, 4a, 5a) may have been related to day-to-day variations in the recorded sEMG signal at each muscle and possibly subtle changes in recording conditions. Sources of variability can compromise the reliability of the under-test metric if the metric is highly sensitive to such changes (which would challenge translational practicality and scaling up the method out of research labs). This paper shows that the topographical measures from muscle networks have less sensitivity to such sources of variations and this can be due to the fact that a muscle network instead of measuring activations at the node level, would calculate higher abstract information regarding the hidden synergistic patterns of functional integration and segregation of activities, enhancing reliability and reducing the effect of stochastic variations at the node level. The presented Spearman-Brown prophecy results provide further insight into the hypothetical required number of measurements to statistically reach excellent between-session reliability. The results show that the network metrics could reach excellent reliability more efficiently than the spectrotemporal metrics with a much less number of sessions consistently across tasks and muscles (Tables V and VI).

## Limitation Statement

This study was limited by the number of tasks and muscles studied and does not provide evidence for upper-limb assessments. Further research is needed to expand on the applications of functional muscle network reliability analysis to other tasks, other limbs, and other muscles.

## V. Conclusion

Functional muscle network analysis is an emerging technique for evaluating the neural control of human motion, it allows to understand the integration and segregation of neural drive distributed from the central nervous system to various parts of the peripheral system. In this study, for the first time, we show that functional muscle network analysis exhibits excellent between- and within-session reliability for submaximal dynamic tasks in the lower limb. This high level of reliability was not observed when using traditional spectrotemporal features. These results emphasize the significance of muscle networks in evaluating functional tasks that involve complex coordination of multiple muscles. Furthermore, the high sensitivity of muscle network to neurophysiological and functional changes recently shown in our work [15], [16], [17], [18] in combination with the high reliability observed in this study, endorses the use of topographical measures of muscle network as a comprehensive neurophysiological biomarker. Given the high clinical relevance of the tasks chosen in this study, the strong reliability across sessions suggests a potential for use in rehabilitation contexts.

## VI. Acknowledgement

The authors would like to acknowledge and thank the help and input from Dr. Yahya Shirazi.

